# Sticking together: Independent evolution of biofilm formation in different species of staphylococci has occurred multiple times via different pathways

**DOI:** 10.1101/2024.03.01.582901

**Authors:** Lisa Crossman, Leanne Sims, Rachael Dean, Heather Felgate, Teresa Diaz Calvo, Claire Hill, Iain McNamara, Mark A Webber, John Wain

**Affiliations:** Quadram Institute Bioscience, Norwich, UK; University of East Anglia, School of Biological Sciences, Norwich, UK; SequenceAnalysis.co.uk, Norwich, UK; University of East Anglia, School of Medicine, Norwich, UK; Norfolk & Norwich University Hospital, Norwich, UK

**Keywords:** Prosthetic joint infection, machine learning, protein domains

## Abstract

Various species of staphylococci cause a wide range of infections, including implant-associated infections which are often difficult to treat due to the presence of biofilms. Whilst some proteins involved in biofilm formation are known, the differences in biofilm production between staphylococcal species remains understudied. Currently biofilm formation by *Staphylococcus aureus* is better understood than for other members of the genus as more research effort has focused on this species.

We assembled a panel of 385 non-*aureus Staphylococcus* isolates of 19 species from prosthetic joint infection as well as other clinical sources and reference strains. We assessed the biofilm forming ability of all strains using a high-throughput crystal violet assay. This identified distinct biofilm formation categories and we then compared the prevalence of Pfam domains and identified those which distinguished the categories as well as using machine learning to identify amino acid 20-mers linked to biofilm formation.

This identified some domains within proteins already positively linked to biofilm formation but we also identified important domains not previously linked to biofilm formation. RT-qPCR confirmed the expression of selected genes predicted to encode important domains within biofilms in *Staphylococcus epidermidis*.

The prevalence and distribution of biofilm associated domains showed a link to phylogeny, suggesting different *Staphylococcus* species have independently evolved different mechanisms of biofilm production.

This work has identified different routes to biofilm formation in diverse species of *Staphylococcus* as well as suggesting independent evolution of biofilm has occurred multiple times across the genus. Understanding the mechanisms of biofilm formation in any given species is likely to require detailed study of relevant strains and the ability to generalise across the genus may be limited.

## Introduction

The genus *Staphylococcus* contains over 60 known species (https://lpsn.dsmz.de/genus/staphylococcus) but clinically has been traditionally split into two groups based on organisms being coagulase negative (CoNS) or coagulase positive. This has proved a pragmatic way to quickly aid the putative identification of *S. aureus* (a coagulase positive species) in clinical microbiology as *S. aureus* infections are common and this species demonstrates high pathogenic potential and antimicrobial resistance (e.g. Methicillin resistant *S. aureus* ‘MRSA’) [1, 2].

However, this distinction is phylogenetically simplistic with most, but not all, non-aureus species of *Staphylococcus* being coagulase negative. The pathogenic potential of CoNS has tended to be underappreciated partly as they are common commensals of the human skin. This has resulted in their isolation often being reported as contamination, not linked to a primary infection. It is now appreciated that many CoNS are important and are often opportunistic pathogens capable of causing a wide range of infections which can be fatal [3].

One area where CoNS are a major cause of infection is in prosthetic joint infection (PJI) which is increasingly common in ageing populations. For example, over a million people in the UK have a replacement joint with >160,000 primary replacements performed annually. Infection is a major complication often necessitating revision with >9500 revisions required per year in the UK [4]. With the average hip revision costing approximately £50,000 [5] this represents an enormous cost in modern healthcare. Infection is one of the leading causes of removal and replacement of implants which itself carries a higher rate of infection than the initial procedure of up to 16% [6]. CoNS are the most isolated pathogens from PJI patients in Europe [7] demonstrating their importance as a major clinical problem.

Specifically diagnosing infection of a joint is a major challenge. Joints can become inflamed or loosened due to non-infective reasons (e.g. gout or aseptic loosening), the management of which may not require the extensive surgery needed to replace an infected implant, nor the use of antibiotics. CoNS as a leading cause of PJI makes diagnosing infection more complex; recovery of these organisms from a sample is often not specific, as it is unclear if the organism was present in the joint or picked up from layers of the skin during sampling.

Many bacterial infections involve formation of a biofilm – a community of aggregated cells which are typically highly resistant to host defences and antibiotics. Infection of indwelling devices by staphylococci is associated with the formation of biofilm on either native tissue (bone, cartilage) or implanted biomaterial (e.g. catheters and orthopaedic devices) [8]. Formation of a biofilm is an important factor in the pathogenesis of PJI, and CoNS are often associated with chronic biofilm infections on the indwelling joint, [3, 9] Given the importance of biofilms in infection, understanding how they form is crucial for both diagnosis and management of infection. Formation of a biofilm by *Staphylococcus spp*. is complex but there are some generally agreed stages comprising (i) initial attachment to a surface, (ii) production of ‘microbial surface components recognizing adhesive matrix molecules’ (MSCRAMMs) allowing tight adhesion to that surface [10], (iii) proliferation and maturation of the biofilm with expansion of biomass and matrix production (iv) expression of disruption factors to allow detachment of cells to facilitate further colonisation of new surfaces.

Biofilm formation by *S. aureus* has been studied extensively [11] and data from *S. aureus* has been extrapolated to other *Staphylococcus* species despite the large-scale genetic differences between the organisms. The best described system for biofilm matrix production in staphylococci may be the polysaccharide intercellular adhesin (PIA) encoded by the *ica* locus comprising *icaABCD* genes [12]. This polysaccharide is also known as poly-N-acetylglucosamine (PNAG), which shares compositional similarity with chitin, another N-acetylglucosamine homopolymer. Whilst PIA production has been clearly shown to influence biofilm formation various surface proteins such as fibronectin-binding proteins (FnBPs), staphylococcal protein A (SpA), and biofilm-associated protein (Bap) have also been shown to play important roles in biofilm formation. These are characterised by their large size with repetitive domains containing multiple “sticky” adhesins [12].

Whilst PIA can clearly impact biofilm formation, several studies have documented the ability to efficiently form biofilms by *ica* negative strains of both *S. aureus* and *S. epidermidis* [13] including the ability to switch from a polysaccharidic to proteinaceous biofilm in an *icaC* mutant of *S. epidermidis* [14]. Microbial surface components recognizing adhesive matrix molecules (MSCRAMMs) appear particularly important in *S. epidermidis* [15]. For example, SdrGFH allows an attachment mechanism known as dock, lock and latch [16]. Other proteins which have been implicated in biofilm formation in *S. epidermidis* include members of the G5 repeat family such as SasG, Aap and Bhp (equivalent of Bap in *S. aureus*), Bbp (bone sialoprotein-binding protein) and FnBPs (fibronectin binding proteins) [16]. These observations suggest that CoNS can form biofilms using varying molecular machinery and that the genes involved in biofilm formation differ between species and strains.

We have recently assembled and sequenced a large panel of CoNS representing many different species [17]. In this study we aimed to link genotype to biofilm formation in these isolates, and to apply machine learning to identify links between the level of biofilm formation and protein sequences in the isolates tested. This allowed us to identify distinct mechanisms underpinning biofilm formation have evolved in different groups of staphylococci.

## Results

### Strain collection and biofilm formation

A collection of 385 CoNS from clinical samples, healthy human volunteers, animals and type cultures, were genome sequenced [17]. These isolates represented 19 species (Supplementary table 1). These included *S. sciuri* (3) and *S. vitulinus* (3) which have been proposed to be reclassified to the *Mammaliicoccus* genus [18] although most recent research based on analysis of conserved protein content suggests they should remain within the *Staphylococcus* genus [19] so we retained these isolates in our analyses.

**Table 1.** Proteins from *S. epidermidis* with previously reported involvement in biofilm formation with Pfam domains identified here as significantly associated with biofilm formation.

| Protein | Proposed function | Domains identified as significant |
| --- | --- | --- |
| Bhp | Attachment to abiotic surfaces | Y_Y_Y; HYR; PKD; Bre5 |
| Ebh | Fibrinonectin binding | Rib; |
| SdrF | Adhesion to collagen | [Y_Y_Y] |
| SdrG | Adhesion to abiotic surfaces | Sdr_G_C_C |
| Aap | Adhesion to abiotic surfaces | G5 |
| IcaA | Synthesis of PIA | Chitin_synth_2 |

A population structure based on alignments of concatenated amino acid sequences of 16 conserved ribosomal proteins [20] was used to generate a phylogenetic tree of the isolates. This resolved the population into 15 main clusters, as determined by HeirBAPS (Figure 1). Most clinical isolates from cases of prosthetic joint infection were from a cluster predominantly containing strains of *S. epidermidis*.

**Figure 1.**
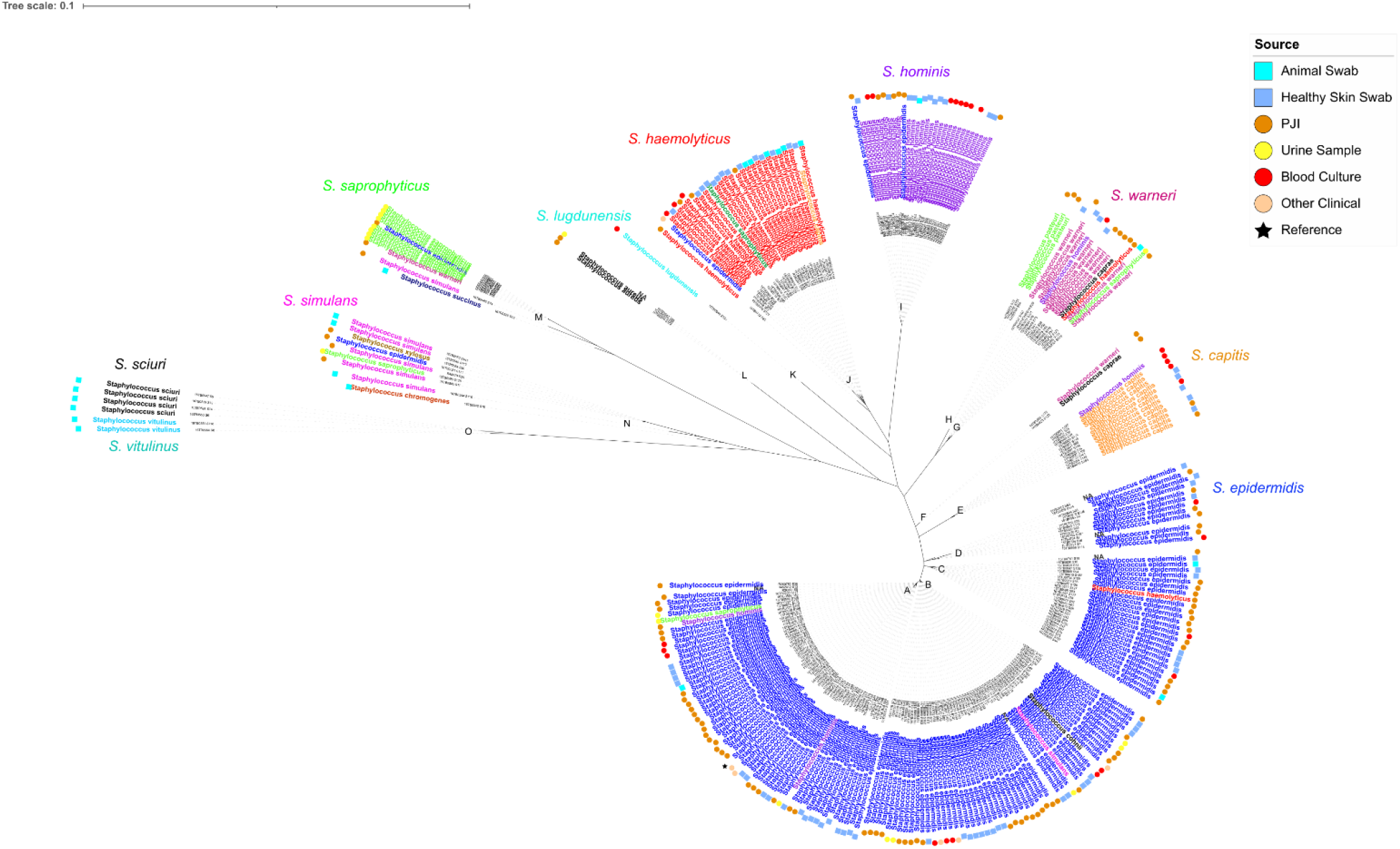
Unrooted maximum likelihood phylogenetic tree based on alignment of concatenated ribosomal protein sequences from all strains. Coloured names indicate different species assignments from MALDI-TOF. Coloured circles on the outer rings indicate sources of isolates.

Biofilm formation by all isolates was determined using a high throughput crystal violet assay. Analysis of the data identified four clearly separated levels of biomass production (Figure 2, Supplementary table 1) by different strains. Each strain was assigned a biofilm level (1-4, with 1 being the lowest and 4 being the highest based on biomass staining). These levels reflected low/no biofilm producers (46.3% of isolates, 161/348, A_595_ <1.15), moderate biofilm producers (22.1% of isolates, 77/348, A_595_ 1.15-2.50), strong biofilm producers (12.1% of isolates, 42/348, A_595_ 2.50-3.85), and very strong biofilm production (19.5% of isolates, 68/348, A_595_ ≥3.85).

**Figure 2.**
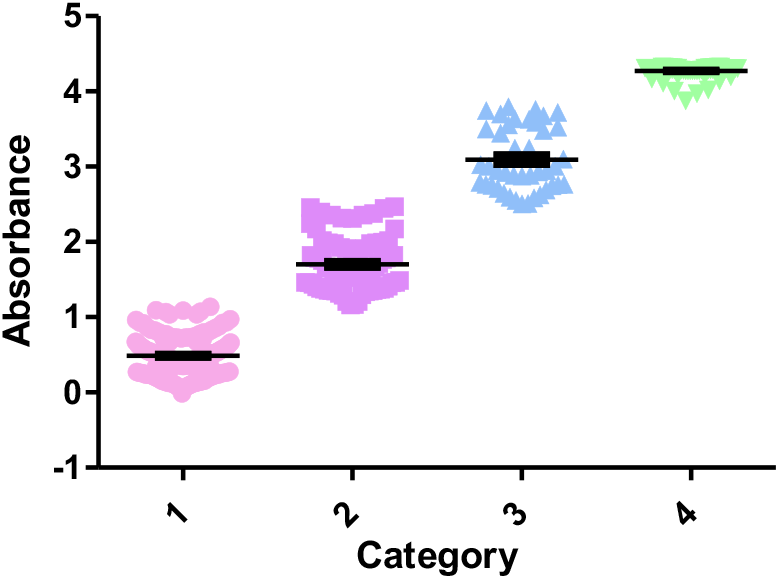
Average biomass production by all isolates based on staining with crystal violet and measured at an absorbance of 595 nm. Each spot represents average biomass for an individual strain and is based on data from four independent replicates.

### Identification of protein family domains and correlation with biofilm formation

All the genomes of the isolates were annotated, and the protein domains present determined by analysis of the Pfam database. Pfam is a large and comprehensive database of protein families where each entry represents a group of related protein sequences sharing a common evolutionary origin. Pfam domains are defined as conserved regions within a protein sequence that are responsible for a specific function or structural motif.

We counted the number of Pfam domains across all predicted proteins in the sequenced isolates. Domain counts were collected and normalised as described previously [21] and the distribution of Pfam domain counts between groups with different biofilm forming capacity was analysed using DESeq2. Significant differences (p-adj. <0.05) were found to be present in the collection.

Domains were identified as being positively or negatively correlated with biofilm formation and we identified 21 domains with significantly different counts across the dataset (according to adjusted P values, Figure 3 and Supplementary Table 2). Of these, 14 were positively linked to biofilm formation, whereas 7 were negatively linked. The positively linked domains included domains found in staphylococcal proteins previously linked to biofilm formation. Key proteins included IcaA (polysaccharide intercellular adhesin synthase, containing domain Chitin_synth_2), SraP (serine-rich adhesin for platelets, containing the He_PIG domain), SdrG (surface-associated fibrinogen binding protein, containing domain SdrG_C_C) and Aap (accumulation-associated protein, containing domain G5). Domains not previously linked to biofilm-specific proteins but positively linked to biofilm formation in this study included the His_biosynth domain and Apc3. The His_biosynth domain is found in histidine biosynthetic proteins, whereas Apc3 is found in TPR-domain containing proteins. These represent interesting candidates as novel proteins involved in biofilm formation. Conversely, domains from proteins which play a role in thiamine synthesis (Thi4), 4Fe-4S cluster formation (Fer4) and hypothetical membrane domains of unknown function (EamA) were found to be negatively linked to biofilm formation.

**Table 2.**
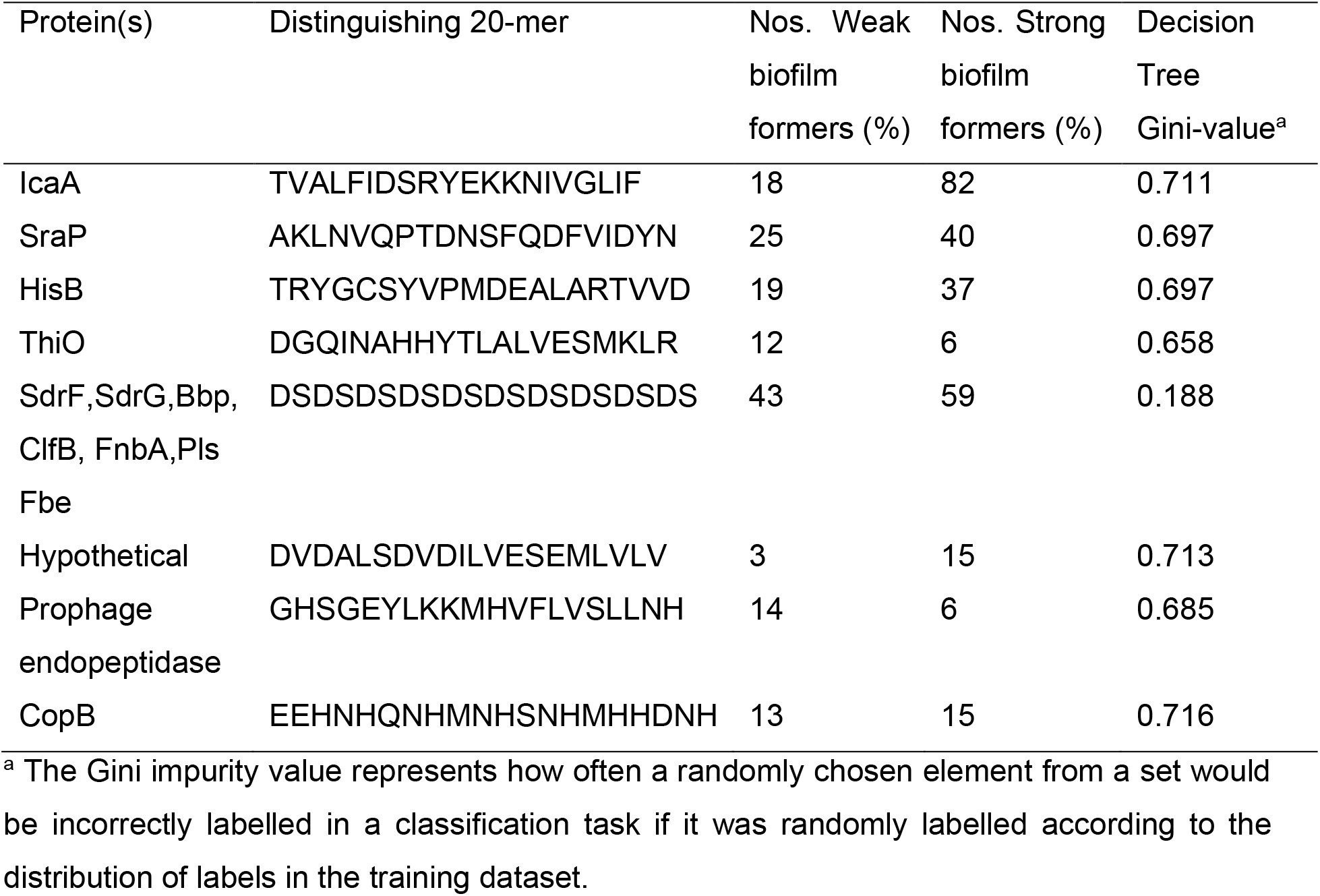
Amino acid sequences of 20-mers identified to be important in non-*aureus* staphylococcal biofilm formation by machine learning. Findings relate to the decision tree in Supplementary Figure 1.

**Figure 3:**
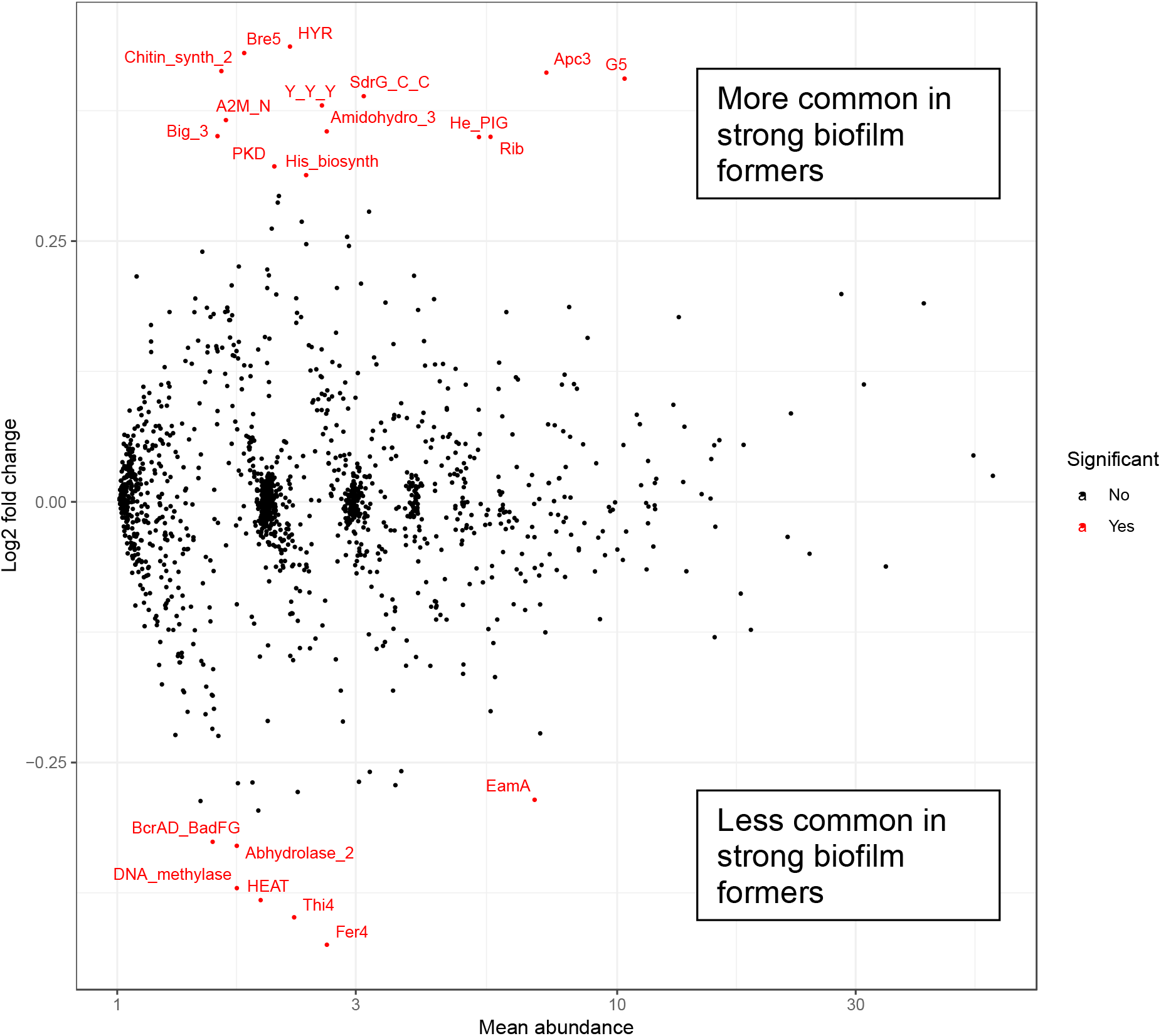
Differential presence of Pfam domains in high vs low biofilm forming strains of non-*aureus* staphylococci, calculated using DESeq2. Domains calculated to be significantly differentially abundant (P-adj <0.05) are highlighted in red.

A PCA analysis of the domains identified as important to biofilm formation using Kruskal-Wallis significance testing clearly showed that five different clusters were present and that these groups related to the phylogeny of the strains (Figure 4). Some of these clusters of domains were associated with a group including multiple species (e.g. a set of domains were clustered in group 1 observed in *S. haemolyticus, capitis, saprophyticus* and *warneri* strains) whereas some were species specific (e.g. groups only seen for *S. hominis* and *simulans*). Different clusters were associated with either stronger or weaker biofilm formation. Diverse strains of *S. epidermidis* were found in more than one cluster (groups 2 and 3 in Figure 4), various proteins have been previously documented as having a role in biofilm formation in *S. epidermidis* (Foster et al., 2020) and there is a lot of genetic diversity present across isolates of this species. Our results suggest that different strains of *S. epidermidis* have acquired different mechanisms of biofilm formation. To explore this further we identified Pfam domains in proteins associated with biofilm formation that were relevant to *S. epidermidis* in the analysis. The results are shown in Table 1.

**Figure 4:**
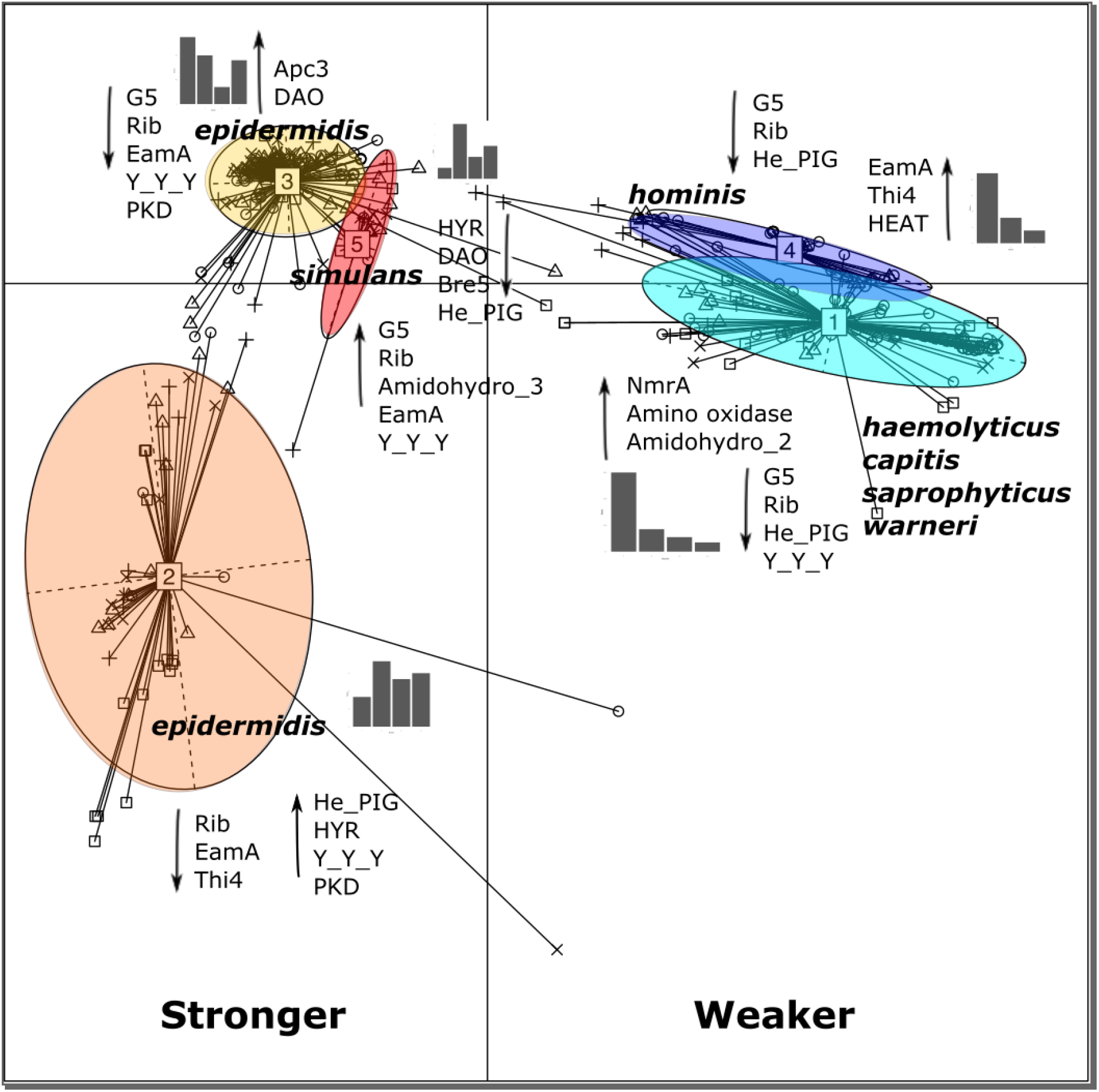
Normalized counts of Pfam domains with significantly different abundance were plotted using principal component analysis (PCA) using a Jensen-Shannon Divergence distance function, with partitioning around medoids (PAM) clustering and between class analysis to identify the principal components. The resulting plot indicated 5 groups were present and these were compared with metadata for species name, biofilm formation and domain count information. Bar graphs indicate the numbers of strains in each cluster displaying each level of biofilm (1,2,3,4). Higher Pfam domain counts are indicated by up-arrows and lower by down-arrows. Radial lines and symbols indicate individual strains with level of biofilm (0(unknown) – square, 1-circle, 2 – triangle point up, 3 - plus, 4 - cross)

We repeated the Kruskal-Wallis significance testing after splitting the strains into phylogenetic groups (rather than on groups based on biofilm forming ability) to gain greater insight into the mechanisms used by separate groups of *Staphylococcus*. The group of greatest interest was the cluster of mostly *Staphylococcus epidermidis*, where the majority of PJI clinical isolates were grouped (Figure 1, groups A-D). Low levels of Pfam domains associated with Ica proteins were present in this group across all levels of biofilm formation. Ica proteins were however found in higher levels in the highest biofilm-forming category of a group containing mostly *S. simulans* (Figure 1, group N). Conversely, the domain Big_3 (found in bacterial surface proteins with immunoglobulin-like folds) was identified in high biofilm formers from one of the *epidermidis* clusters (biofilm category 3, Figure 1 groups A-D), as well as very high biofilm formers of a cluster of *hominis* strains (Figure 1, group I), and *S. chromogenes* isolates, which were again all very strong biofilm formers (Figure 1, group N). These data again demonstrate distinct biofilm formation mechanisms have evolved between different phylogenetic groups.

### Machine learning decision trees

To further identify protein signatures linked with different abilities to form biofilm we used a machine learning approach. We generated a subset of 180 genome sequences to produce two alternative input training datasets, one with the proteins containing the significant Pfam domains only, and another containing all the predicted proteins across the genomes. This subset was chosen to provide balance between species and degrees of biofilm formation and strains were randomly chosen to fulfil these criteria. The number of genomes represented in each category of biofilm formation (1-4) were 69:53:18:40, respectively.

A machine learning workflow was built for the dataset by initially splitting the predicted protein sequences into 20-mer ‘word’ sequences (non-overlapping). The strain data were aggregated by biofilm metadata type into sets of words for each biofilm level: 1, 2, 3 and 4, with those labels. For regression, the labels were provided as numbers, whereas for classification, labels were supplied as character data. Using the labels in these means in regression, the biofilm levels were taken as a continuous dataset where 1 was regarded as the lowest and 4 the highest form of biofilm type. For classification, the biofilm levels were presented as character labels, meaning each biofilm level was represented as a unique entity.

The vectorized dataset (see methods) was fitted with either a Decision Tree Regressor or a Decision Tree Classifier with a maximum depth of 15. Trees were visualised, and model accuracy determined against the test data (Supplementary Figure 1). For the Pfam dataset, the model accuracy reached on a single decision tree was 70% for the best fit regressor but was not improved by a random forest approach. For the best fit classifier, 68% was reached on a single decision tree. The model accuracy at predicting strong or weak biofilm formation was increased by using Extratrees to 75%, suggesting that passing the label as a classification task was appropriate. This is in accordance with the Pfam PCA plot (Figure 4), which indicated that alternative means of biofilm formation are evident in different groups of CoNS. Random Forests were also tested with a maximum accuracy of 74%. The accuracy of predicting the biofilm category of a single isolate (1-4) remained at 63%, a greater accuracy was probably not possible due to multiple factors including the inclusion of different species carrying out alternative strategies, inherent variation in the crystal violet assay and overfitting towards the lower leaves, particularly in distinguishing between biofilm levels 1 and 2. For the whole genome dataset, an accuracy of 67% could be reached with regression, whilst 77% accuracy could be reached with classification boosted by ExtraTrees.

Some of the 20-mers identified were in themselves capable of distinguishing between strong and weak biofilm formation, decision tree depths represent level at which the sequence was determined to be differentiating between weak and strong biofilm formers (Supplementary Figure 1). For example, despite IcaA being present in both strong and weak biofilm forming strains, the exact 20mer ‘TVALFIDSRYEKKNIVGLIF’ (depth 1 in the Pfam decision tree) found within IcaA could be used to distinguish the groups. Of the strains containing this 20-mer, 82% were strong biofilm formers (categories 3 and 4), whereas 73% of the strains containing IcaA without this 20-mer were weak biofilm formers (categories 1 and 2). This appears to be due to variations, rather than truncations within the protein sequences, as 368 IcaA sequences from various isolates had a mean length of 407.02 amino acids, whereas 148 IcaA sequences without the 20-mer had a mean length of 407.95 amino acids. This suggests that small variations within the sequence of IcaA can have large impacts on biofilm formation ability.

A further 20-mer was identified within the sequence of SraP (Serine rich adhesin for platelets) ‘AKLNVQPTDNSFQDFVIDYN’. This was present in 40% of strong biofilm formers and only 25% of weak biofilm formers with 48% of strong (category 4) strains having this sequence. The SraP protein was identified in 6 alternative nodes with depths 6,7,8,9,10 & 14 and at depths 2 and 4 in the whole genome decision tree. Similarly a 20-mer was identified within HisB ‘TRYGCSYVPMDEALARTVVD’ that was associated with strong biofilm formation with 37% of category 4 strains possessing this sequence compared with 19% of category 1 strains. A fourth sequence from ThiO (glycine oxidase) was correlated with weak biofilm formation ‘DGQINAHHYTLALVESMKLR’ and was present in 12% of weak biofilm formers versus only 6% of strong biofilm formers. ThiO was found in 2 alternative nodes at depths 4 and 8 in the Pfam tree. One 20-mer sequence motif ‘DSDSDSDSDSDSDSDSDSDS’ was identified in several different proteins, including SdrF, SdrG (Serine-aspartate repeat-proteins), Bbp (bone sialoprotein-binding precursor), ClfB (clumping factor B), FnbA (fibronectin binding protein A), Pls (Surface protein precursor) and Fbe (fibrinogen binding precursor) with 59% of strong (category 4) biofilm formers having this motif compared to 43% of weak isolates. The Sdr proteins were identified in 2 alternative nodes at depths 5, 8, 16 and 17. Another set of important 20-mers were also identified using the same method on the whole genome sequences of the collection. These included a 20-mer identified in a large hypothetical hydrophobic protein belonging to the serine-rich repeat proteins, ‘DVDALSDVDILVESEMLVLV’ (depth 1 in the whole genome decision tree), this sequence was over-represented in strong vs weak biofilm forming strains (15 vs 3, respectively). A 20-mer in prophage endopeptidase sequences, ‘GHSGEYLKKMHVFLVSLLNH’ (depth 2 in the whole genome tree), was found to be under-represented in strong vs weak biofilm forming strains (16 vs 4, respectively). Finally, a 20-mer was identified in CopB, ‘EEHNHQNHMNHSNHMHHDNH’ (depth 4 in the whole genome tree). This was found to be similarly present in strong and weak biofilm formers (15 vs 13, respectively), but could better distinguish between category 3 and category 4 biofilm formers (14.5 vs 0.8 %, respectively). Lower decision tree rank depths may represent some overfitting with respect to the two categories of low biofilm formers 1 and 2, but do suggest a high level of discrimination.

### Validation of predicted contributions of genes in a model of PJI

A sub-selection of genome assemblies of *S. epidermidis* isolated from cases of PJI and belonging to the largest phylogenetic cluster (Figure 1, groups A-D) was used to check for the presence of genes containing domains identified by machine learning as contributing to biofilm formation. Of the genes identified in this study, five were present in all PJI isolates checked, therefore carried forward for further investigation into their expression profiles in *Staphylococcus epidermidis* biofilms. These genes were the imidazoleglycerol-phosphate dehydratase-encoding *hisB*, the copper-transporting P-type ATPase-encoding *copB*, a prophage endopeptidase (PE), and a hypothetical hydrophobic protein which the sequence suggests belongs to the serine-rich repeat family, referred to as HH here.

Expression of these genes was then measured in two different isolates. *Staphylococcus epidermidis* RP62A (DSM 28319) is a well-studied model biofilm forming strain that contains the *ica* genes (Figure 1, group A). Conversely, strain 15TB0846 (hereafter referred to as ‘846’) was identified as a very strong biofilm former which did not contain the *ica* operon (Figure 1, group A), suggesting other mechanisms must allow biofilm formation.

Both strains were grown as biofilms and planktonic cultures before RNA extraction, and quantification of gene expression using RT-qPCR. Relative expression in the biofilm compared to planktonic conditions was calculated as a Log_2_ fold using *gyrB* as a reference gene (Figure 5).

**Figure 5.**
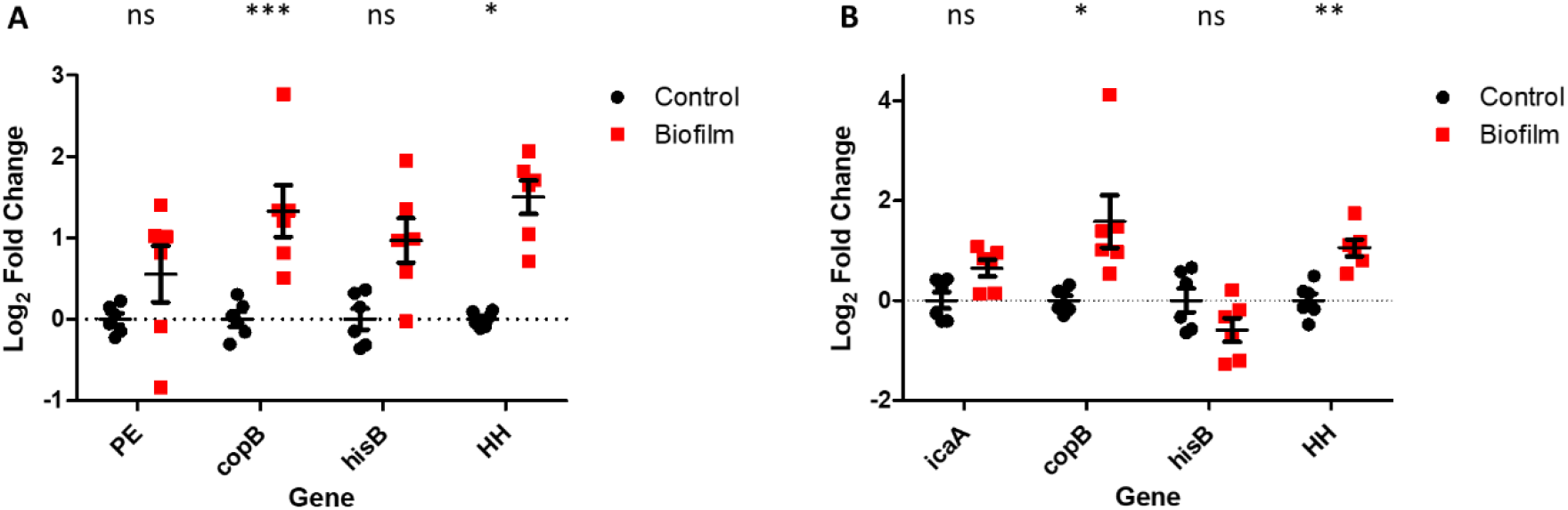
Relative expression of genes associated with biofilm formation by machine learning in *Staphylococcus epidermidis* strains A) 846 and B) RP62A, change in expression in biofilms calculated relative to planktonic cells, relative to gyrB as a housekeeping gene. Error bars show SEM (n=6). * = P<0.05, ** = P<0.01, *** = P<0.001.

Expression of all genes identified by machine learning as important for biofilm formation was confirmed and *copB* and HH expression was significantly upregulated in both strains (p<0.05) in biofilms. Expression of *icaA* (only present in *S. epidermidis* RP62A) was also upregulated in biofilm growth as expected.

## Discussion

In this study, we used the relative abundance of Pfam domains to identify proteins involved in biofilm formation. We showed that the Pfam domains: G5, Rib, He_PIG and Y_Y_Y, A2M_N and Bre5 are correlated with a strong biofilm-forming phenotype.

The G5 domain is found in the Aap (accumulation-associated) protein of *Staphylococcus epidermidis*, which has been linked to PIA-independent biofilm formation in the context of PJI [22]. Rib domains are present in fibronectin binding protein Ebh, and the biofilm-associated protein Bap, both of which have also been demonstrated to play a role in biofilm formation [13]. The He_PIG domain is present in the serine-rich adhesin for platelets (SraP) protein characterised in *Staphylococcus aureus* [23], and Y_Y_Y, A2M_N and Bre5 are found in Bhp, a homologue of the aforementioned Bap; Y_Y_Y is also present in SdrF, which plays a role in adhesion to abiotic surfaces [24]. This demonstrates the wide range of proteins involved in biofilm formation in coagulase-negative *Staphylococcus* isolates.

Most domains identified here were part of proteins known to act as adhesins, although not all. For example, His_biosynth and Apc3 were associated with biofilm formation but are not linked directly to their presence in adhesion proteins.

The HisB protein is an imidazole glycerol phosphate dehydrogenase involved in histidine biosynthesis - a mutation in *hisB* reduces biofilm forming ability in *Staphylococcus xylosus* [25]. Histidine is commonly found in membrane proteins [26] which play an important role in biofilm formation [24]. This could explain the presence of, histidine, lysine and arginine at the surface of the cell and the link between biofilm formation and the biosynthesis of positively charged amino acids.

The Apc3 domain is present in the amino acid sequences of SERP1184, SERP1033 and SERP0431 in *Staphylococcus epidermidis* RP62A. These genes all encode for TPR-repeat containing proteins, a motif which plays an indirect role in adhesion to host cells and biofilm formation through their role in type IV pilus biogenesis [27].

Our PCA clusters of the Pfam domains associated with strong biofilm formation suggested an association between *Staphylococcus* species and biofilm mechanism (Figure 4, left hand panel). The Ica proteins are responsible for the biosynthesis of polysaccharide intracellular adhesin (PIA), a key biofilm component in many strains of *Staphylococcus epidermidis* [13] and *Staphylococcus hominis* [28], whereas *Staphylococcus aureus* biofilms are more often protein-dependent [13]. PIA also plays a relatively small role in biofilm formation of *S. haemolyticus*, where eDNA and protein components were more important [29]. A study of *Staphylococcus* species isolated from bovine mastitis found the *icaA* gene to be present in a range of CoNS, including *Staphylococcus chromogenes, sciuri* and *xylosus*, and *S. aureus* isolates, however it was not found in *Staphylococcus epidermidis* isolates, demonstrating the ability of *S. epidermidis* to use different methods for biofilm formation [30].

A further range of proteins, linked to high levels of biofilm formation, were identified when we applied machine learning to the biofilm output associated with draft genome assemblies, separating the protein sequences into amino acid 20-mers and building a decision tree to differentiate between levels of biofilm formation. We focussed on a subset of genes identified using this method based on their presence in staphylococcal isolates from prosthetic joint infection, where biofilm formation has a severe effect on the treatment options and prognosis [9]. The list generated included IcaA (and other Ica proteins), HisB, CopB, prophage endopeptidase, and a hypothetical hydrophobic protein. HisB and IcaA are both well-known and serve as a positive control for our method.

The hypothetical hydrophobic protein 20-mer was identified in proteins belonging to the serine-rich-repeat family of adhesins, containing an N-terminal signal peptide, short serine-rich repeat (SRR) domain, ligand binding domain, longer SRR domain and C-terminal LPXTG motif for cell wall anchoring, playing roles in both biofilm formation and virulence [31]. It is of note that many draft assemblies did not have this SRR protein accurately annotated (due to the repeat regions complicating assembly from short-read sequencing data) and so further discussion is warranted here. The best characterised of these is the serine-rich adhesin for platelets (SraP) in *Staphylococcus aureus* [32], which has a homology of 56% at the amino acid level to the protein of interest from *Staphylococcus epidermidis* 846. Another protein from this family, UafB from *S. saprophyticus*, mediates binding to fibrinogen, fibronectin and human uroepithelial cells [33], and is 46% identical to the *S. epidermidis* 846 protein of interest. This demonstrates the ability of a machine learning technique trained on 20-mers (independent of annotation) to identify key proteins of interest without the need for complete genome assemblies.

We also identified two novel proteins involved in staphylococcal biofilm formation, CopB and prophage endopeptidase. CopB is a copper transporting P-type ATPase. To our knowledge, this protein has not been previously linked to biofilm formation in *Staphylococcus*, however a *copB* mutant of the plant pathogen *Xylella fastidiosa* produced higher amounts of biofilm than the wild type [34]. The ability to tolerate high levels of copper (among other metals such as Zn, As and Cd) has been linked to the ability of *S. saprophyticus* to cause infections [35]. The introduction of *copB* and *mco* (also identified here using machine learning) to a naïve clinical isolate of *Staphylococcus aureus* conferred hyper tolerance to copper, which was linked to virulence [36].

The identification of prophage endopeptidase is again not previously linked to biofilm formation, yet phage islands have been linked to invasiveness in a comparative study of colonizing and invasive *Staphylococcus epidermidis* from patients with prosthetic joint infection [37], where 226/299 infection-associated genes mapped to prophage regions.

RT-qPCR was used to confirm selected genes containing novel domains predicted to be important for biofilm formation were expressed in biofilms. Expression of *copB* and the gene encoding the SRR protein were significantly upregulated in biofilms compared to planktonic growth suggesting their importance in biofilm formation.

## Conclusions

Several biofilm forming mechanisms were described in the coagulase-negative staphylococci with a possible link to species (or sub-species) but there was no difference between isolates from PJI and other samples. This is similar to reported findings with *Staphylococcus haemolyticus* [29] but converse to invasive *Staphylococcus epidermidis* isolates from PJI which form larger biofilms and were more likely to contain mobile genetic elements then comparators [37].

The large complement of genes identified here as being linked to biofilm formation within different phylogenetic groups suggest that the ability to form a biofilm is a fundamental part of the biology of staphylococci that has evolved multiple times and is encoded redundantly on the genome. Although no definable combination of genes can predict, or indicate, biofilm ability, limiting the opportunity to develop diagnostic assays, convergent functions (such as adhesion) may be accessible.

Further research is needed to better stratify and so understand the mechanisms of biofilm formation present in different CoNS and to exploit this knowledge to develop new strategies for preventing and treating CoNS infections.

## Methods & Materials

### Quantifying biofilm formation by Staphylococcus isolates

A collection of *Staphylococcus* isolates was obtained from a mixture of swabs of healthy volunteers and clinical samples from the Norfolk and Norwich University Hospital. Isolates were identified to the species level using MALDI-TOF and combined with rare isolates from the National Collection of Type Cultures to yield a total of 385 strains for analysis.

The level of biofilm formation was quantified using a modified version of the crystal violet assay outlined in [38]. Briefly, isolates were streaked onto Columbia blood agar plates and incubated overnight at 37 °C. A sample of 3-5 colonies was used to inoculate 5 mL HHW medium [39], and grown at 37 °C with shaking overnight. These cultures were diluted 1/200 in HHW pre-warmed to 37 °C, and 150 μl aliquots were grown in 96-well plates, with four replicates for each isolate. Positive and negative biofilms were performed on each plate using two strains identified as highest and lowest biofilm formers, 15TB0798 and 15TB0711 respectively. Medium without inoculum was used as a sterility control. The plates were incubated at 37 °C for 24h with 100 rpm shaking. The contents were discarded, and the wells washed with PBS, followed by fixing with 200 μL ethanol for 15 minutes. Excess ethanol was removed, and the plates dried, then the biofilms were stained with 200 μL 2% crystal violet solution. The plate was then washed with water, and the crystal violet resolubilised using 200 μl glacial acetic acid in water. The plates were sealed and vortexed before OD_595_ readings were taken. Measurements were normalised by subtracting the absorbance value from the sterility control wells. Strains were categorised according to the mean of the normalised absorbance readings over four replicates. Average readings below 1.15 were assigned to category 1 (low/no biofilm formation), between 1.15 and 2.50 were assigned to category 2 (moderate biofilm formation), between 2.50 and 3.85 were assigned to category 3 (high biofilm formation), and above 3.85 were assigned to category 4 (very high biofilm formation).

### DNA extraction, sequencing and genome assembly

DNA extraction was performed using the QIAGEN QIAcube. A 1 ml aliquot of overnight culture grown in TSB was harvested by centrifugation, and the cell pellet was resuspended in 400 μL buffer AE containing Reagent DX (100 μL/15mL). Cell suspensions were transferred to 2 mL lysing matrix B tubes (MPBio), followed by bead-beating in a TissueLyser II at 30 Hz for 15 min, turning halfway through. The lysed samples were centrifuged at room temperature (5000 ×*g*, 5 min), before transferring the supernatant into the S block and adding 4 mL RNaseA and using the QIAcube and QIAamp DNA mini kit according to manufacturer’s instructions.

Nextera XT library preparation was performed according to the manufacturer’s protocol, and strains were sequenced using Illumina MiSeq or NextSeq machines in 2 × 150 bp cycles. Genome assembly was performed using SPAdes [40] after pre-processing and read coverage normalisation on any samples with over 70× coverage. Annotation was carried out using Prodigal [41].

### Pfam domain identification and association with biofilm ability

Methods to produce the dataset were followed as described previously [21]. Briefly, draft genome assemblies were input as protein fasta files, using a grouping file to differentiate between the biofilm categories. HMMER3 [42] was run against the Pfam A database [43], and separate Pfam tables were combined. Statistical significance testing between groups was carried out using Kruskal-Wallis testing or DESeq2 [44], as indicated in the relevant text. PCA plots were produced using the ade4 R library [45]. The version of the Pfam database used in this study was 32.0 (2018-08-30).

### Machine learning

The machine learning model was trained with the use of scikit-learn library version 0.23.2. The complete code encoding the protein sequences for whole genome and Pfam domain counts datasets is available on https://github.com/LCrossman. Initially, the dataset was split into 25% test data and 75% training data and vectorized using a term weighting scheme commonly used to represent text documents as the normalized term frequency (number of occurrences) of each term in a document. The vectorized dataset was fitted with either a Decision Tree Regressor or a Decision Tree Classifier with maximum depth of 15. The model accuracy at predicting strong or weak biofilm formation was increased using the ensemble method Extremely Randomized Trees (Extratrees) [46]. Random Forests were also tested, however, the best accuracy at 77% was achieved using biofilm labels as categorical with classification decision trees boosted by the ExtraTrees method.

### Bulk biofilm growth and RNA extraction

*S. epidermidis* RP62A biofilms were grown in 5 mL Mueller-Hinton broth at 25 °C for 72 h, whereas 846 biofilms were grown in HHW [39] at 37 °C for 24 h. Both were grown on discs of steel 316L with 40 rpm orbital shaking. Biofilms were harvested from the discs by washing twice in PBS, before vortexing in 3 mL PBS. The resulting cell suspension was centrifuged to pellet the cells, and the supernatant discarded. Control cultures were grown with 180 rpm shaking and in the absence of discs to prevent biofilm formation. The pellet was immediately resuspended in 100 μL lysis buffer containing 20 mM Tris-HCl (pH 8.0), 2 mM EDTA and 0.5 mg mL^-1^ lysostaphin, followed by incubation at 37 °C for 10 min. After this, the Promega SV Total RNA Isolation Kit was used according to manufacturer’s instructions. RNA quantity and purity was measured using Qubit DNA and RNA quantification, and NanoDrop. RNA quality was assessed using an Agilent 2200 TapeStation, and samples with an RNA Integrity Number above 5 were used for RT-qPCR analysis.

### RT qPCR primers – design and validation

Primers for RT-qPCR (Table 3) were designed using the Primer3 software [47], and checked for potential dimerization and secondary structures using the IDT oligoanalyzer tool. Primers were validated (Supplementary figure 2) using genomic DNA extracted from *Staphylococcus epidermidis* strains RP62A and 846 using the Zymo Quick-DNA miniprep kit according to manufacturer’s instructions, with the addition of an initial 30 min lysis step using 0.5 mg mL^-1^ lysostaphin at 37 °C. RT-qPCR reactions were carried out in 10 μL reactions using the Luna® One-Step Universal RT-qPCR kit (NEB) according to manufacturer’s instructions, with 1 ng target RNA. Expression of genes of interest in biofilm compared to biomarker was calculated as fold-change relative to the reference gene *gyrB* using the 2^ΔΔCt^ method. Significance was calculated using 2-way ANOVA and ΔC_T_ values.

**Table 3.**
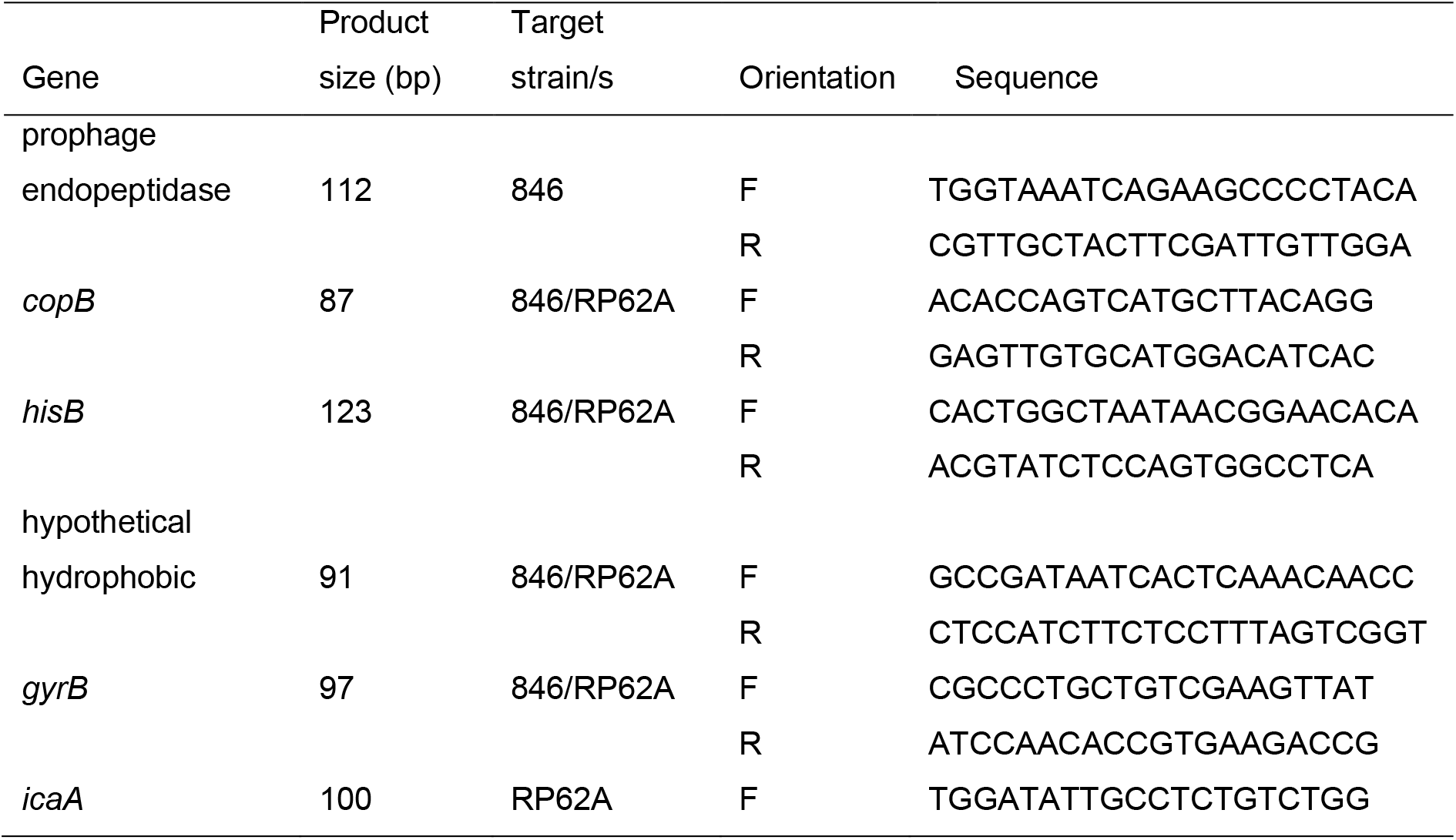
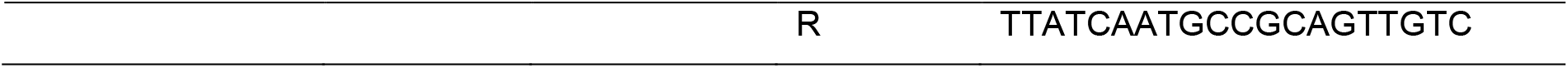
Primers used for RT-qPCR experiments.

## Data Availability

The reads representing the raw genome sequence data used in this project can be accessed as ENA BioProject PRJEB31403. Study accession ERP113963.

Code used for the machine learning work is available from https://github.com/LCrossman

## Acknowledgements

Funding – this work was supported by Heraeus Medical via a grant to IM and JW; and by funding from Action Arthritis to MW, JW and IM. HF was supported by BBSRC grant number BB/T014644/1

## Conflicts of interest

The authors have no conflicts to declare

## Notes

https://www.ebi.ac.uk/ena/browser/view/PRJEB31403

https://github.com/LCrossman

### Competing Interest Statement

The authors have declared no competing interest.

