## Supplementary Information for "Sticking together: Independent evolution of biofilm formation in different species of staphylococci has occurred multiple times via different pathways"

**Supplementary Table 1: Species composition and biofilm phenotype**

| Species (MALDI-TOF designated) | Total isolates | Isolates per biofilm category * | ** | *** | **** |
|---|---|---|---|---|---|
| Unknown *Staphylococcus* | 3 | 2 | 0 | 0 | 1 |
| *Staphylococcus auricularis* | 2 | 2 | 0 | 0 | 0 |
| *Staphylococcus capitis* | 19 | 13 | 2 | 1 | 3 |
| *Staphylococcus caprae* | 2 | 0 | 0 | 1 | 1 |
| *Staphylococcus chromogenes* | 3 | 0 | 0 | 0 | 3 |
| *Staphylococcus cohnii* | 1 | 0 | 0 | 0 | 1 |
| *Staphylococcus epidermidis* | 193 | 62 | 53 | 26 | 52 |
| *Staphylococcus equorum* | 1 | 1 | 0 | 0 | 0 |
| *Staphylococcus haemolyticus* | 44 | 35 | 2 | 5 | 2 |
| *Staphylococcus hominis* | 44 | 31 | 11 | 2 | 0 |
| *Staphylococcus lugdunensis* | 9 | 5 | 2 | 2 | 0 |
| *Staphylococcus pasteuri* | 3 | 2 | 1 | 0 | 0 |
| *Staphylococcus pettenkferi* | 1 | 1 | 0 | 0 | 0 |
| *Staphylococcus saprophyticus* | 22 | 6 | 6 | 5 | 5 |
| *Staphylococcus sciuri* | 5 | 1 | 0 | 3 | 1 |
| *Staphylococcus simulans* | 9 | 0 | 3 | 3 | 3 |
| *Staphylococcus succinus* | 1 | 1 | 0 | 0 | 0 |
| *Staphylococcus vitulinus* | 3 | 2 | 0 | 0 | 1 |
| *Staphylococcus warneri* | 18 | 11 | 5 | 1 | 1 |
| *Staphylococcus xylosus* | 2 | 1 | 0 | 0 | 1 |

**Supplementary Table 2: Top 100 outputs for significant Pfam domains**

| Pfam domain | baseMean[a] | log2 Fold Change[b] | Standard error[c] | stat[d] | P value | Adjusted P value |
|---|---|---|---|---|---|---|
| Apc3 | 7.21779 | 0.41153 | 0.07554 | 5.44807 | 5.09E-08 | 8.65E-05 |
| HYR | 2.21602 | 0.43653 | 0.08514 | 5.12748 | 2.94E-07 | 0.000249 |
| Bre5 | 1.79488 | 0.43034 | 0.08651 | 4.97429 | 6.55E-07 | 0.00036 |
| Fer4 | 2.62659 | -0.42478 | 0.08627 | -4.92390 | 8.48E-07 | 0.00036 |
| G5 | 10.34756 | 0.40584 | 0.08379 | 4.84376 | 1.27E-06 | 0.000433 |
| Chitin_synth_2 | 1.61552 | 0.41303 | 0.08868 | 4.65729 | 3.20E-06 | 0.00088 |
| SdrG_C_C | 3.11266 | 0.38897 | 0.08398 | 4.63153 | 3.63E-06 | 0.00088 |
| Thi4 | 2.25859 | -0.39845 | 0.08790 | -4.53328 | 5.81E-06 | 0.00123 |
| Y_Y_Y | 2.56733 | 0.38010 | 0.08486 | 4.47886 | 7.50E-06 | 0.00142 |
| HEAT | 1.93614 | -0.38199 | 0.08774 | -4.35382 | 1.34E-05 | 0.00227 |
| A2M_N | 1.65023 | 0.36604 | 0.08679 | 4.21737 | 2.47E-05 | 0.00343 |
| DNA_methylase | 1.73501 | -0.37043 | 0.08783 | -4.21749 | 2.47E-05 | 0.00343 |
| He_PIG | 5.29161 | 0.34977 | 0.08321 | 4.20367 | 2.63E-05 | 0.00343 |
| Amidohydro_3 | 2.62556 | 0.35519 | 0.08733 | 4.06720 | 4.76E-05 | 0.00577 |
| Big_3 | 1.58708 | 0.35067 | 0.08710 | 4.02591 | 5.68E-05 | 0.00642 |
| Rib | 5.58124 | 0.34998 | 0.08797 | 3.97847 | 6.94E-05 | 0.00736 |
| EamA | 6.83724 | -0.28577 | 0.07233 | -3.95095 | 7.78E-05 | 0.00777 |
| Abhydrolase_2 | 1.73488 | -0.32991 | 0.08791 | -3.75286 | 0.000175 | 0.0156 |
| PKD | 2.06183 | 0.32163 | 0.08544 | 3.76455 | 0.000167 | 0.0156 |
| BcrAD_BadFG | 1.55211 | -0.32607 | 0.08788 | -3.71053 | 0.000207 | 0.0176 |
| His_biosynth | 2.38711 | 0.31326 | 0.08676 | 3.61075 | 0.000305 | 0.0247 |
| ABG_transport | 1.91528 | -0.29610 | 0.08778 | -3.37313 | 0.000743 | 0.0574 |
| Gram_pos_anchor | 3.18956 | 0.27819 | 0.08358 | 3.32824 | 0.000874 | 0.0639 |
| HTH_Mga | 2.10600 | 0.29325 | 0.08835 | 3.31905 | 0.000903 | 0.0639 |
| DNA_pol3_gamma3 | 1.46783 | -0.28693 | 0.08774 | -3.27009 | 0.00108 | 0.0702 |
| KorB | 2.09479 | 0.28689 | 0.08767 | 3.27233 | 0.00107 | 0.0702 |
| Amino_oxidase | 3.04457 | -0.26847 | 0.08466 | -3.17133 | 0.00152 | 0.0853 |
| Collagen | 2.88354 | 0.25403 | 0.08026 | 3.16511 | 0.00155 | 0.0853 |
| HI0933_like | 13.28017 | 0.17711 | 0.05532 | 3.20164 | 0.00137 | 0.0853 |
| MarR | 7.01608 | -0.22202 | 0.07058 | -3.14553 | 0.00166 | 0.0853 |
| NAD_binding_4 | 3.60155 | -0.27173 | 0.08621 | -3.15176 | 0.00162 | 0.0853 |
| NmrA | 3.70011 | -0.25834 | 0.08196 | -3.15192 | 0.00162 | 0.0853 |
| Sulfate_transp | 2.29662 | -0.27828 | 0.08733 | -3.18667 | 0.00144 | 0.0853 |
| Y1_Tnp | 2.33946 | 0.26859 | 0.08675 | 3.09606 | 0.00196 | 0.0971 |
| Zeta_toxin | 1.86479 | -0.26914 | 0.08710 | -3.09004 | 0.002 | 0.0971 |
| Flavodoxin_2 | 3.20046 | -0.25899 | 0.08410 | -3.07937 | 0.00207 | 0.0978 |
| Amidohydro_2 | 1.74362 | -0.26987 | 0.08802 | -3.06593 | 0.00217 | 0.0996 |
| Str_synth | 2.03754 | 0.26194 | 0.08782 | 2.98256 | 0.00286 | 0.128 |
| Gly_radical | 2.38824 | 0.24705 | 0.08674 | 2.84814 | 0.0044 | 0.191 |
| Aminotran_5 | 8.01658 | 0.18675 | 0.06601 | 2.82905 | 0.00467 | 0.194 |
| DAO | 31.15039 | 0.11248 | 0.03978 | 2.82761 | 0.00469 | 0.194 |
| IncA | 2.90806 | 0.24535 | 0.08734 | 2.80902 | 0.00497 | 0.201 |
| GTF2I | 1.48010 | 0.23979 | 0.08649 | 2.77247 | 0.00556 | 0.22 |

| | | | | | | |
|---|---|---|---|---|---|---|
| Lipoprotein_Ltp | 3.92379 | 0.21675 | 0.08056 | 2.69071 | 0.00713 | 0.275 |
| FctA | 1.59361 | -0.22450 | 0.08744 | -2.56740 | 0.0102 | 0.382 |
| Glyoxalase | 5.58347 | -0.20082 | 0.07877 | -2.54955 | 0.0108 | 0.382 |
| Ndr | 1.75089 | 0.22564 | 0.08845 | 2.55110 | 0.0107 | 0.382 |
| Z1 | 1.30747 | -0.22362 | 0.08747 | -2.55645 | 0.0106 | 0.382 |
| Peptidase_A24 | 1.99603 | 0.22275 | 0.08794 | 2.53301 | 0.0113 | 0.392 |
| Gate | 6.00348 | 0.18194 | 0.07222 | 2.51936 | 0.0118 | 0.399 |
| ACT | 4.30250 | 0.19432 | 0.07885 | 2.46453 | 0.0137 | 0.409 |
| FMN_red | 2.82824 | -0.21074 | 0.08526 | -2.47160 | 0.0135 | 0.409 |
| Hexapep | 15.66674 | -0.12986 | 0.05269 | -2.46472 | 0.0137 | 0.409 |
| Methyltransf_4 | 3.07418 | 0.20919 | 0.08374 | 2.49811 | 0.0125 | 0.409 |
| Mob_Pre | 2.01137 | 0.21718 | 0.08782 | 2.47293 | 0.0134 | 0.409 |
| NuiA | 1.09365 | 0.21611 | 0.08713 | 2.48049 | 0.0131 | 0.409 |
| Patatin | 1.55021 | -0.21770 | 0.08804 | -2.47284 | 0.0134 | 0.409 |
| YSIRK_signal | 8.72584 | 0.15715 | 0.06458 | 2.43359 | 0.015 | 0.438 |
| AlaDh_PNT_C | 2.75057 | 0.20508 | 0.08522 | 2.40650 | 0.0161 | 0.464 |
| HxlR | 2.00005 | -0.21017 | 0.08773 | -2.39547 | 0.0166 | 0.47 |
| Acetyltransf_1 | 18.50956 | -0.12277 | 0.05301 | -2.31624 | 0.0205 | 0.527 |
| DM13 | 1.99580 | 0.20496 | 0.08787 | 2.33256 | 0.0197 | 0.527 |
| GA | 28.07883 | 0.19923 | 0.08599 | 2.31700 | 0.0205 | 0.527 |
| Ldh_1_C | 3.44001 | 0.19102 | 0.08218 | 2.32427 | 0.0201 | 0.527 |
| Mac | 1.50218 | -0.20371 | 0.08793 | -2.31661 | 0.0205 | 0.527 |
| PRA.CH | 1.69433 | 0.20752 | 0.08850 | 2.34474 | 0.019 | 0.527 |
| TMP.TENI | 3.99749 | 0.18397 | 0.07959 | 2.31150 | 0.0208 | 0.527 |
| PPR | 1.38293 | -0.20140 | 0.08741 | -2.30410 | 0.0212 | 0.53 |
| FIVAR | 41.04384 | 0.19031 | 0.08351 | 2.27887 | 0.0227 | 0.558 |
| Abi | 5.69711 | -0.16809 | 0.07416 | -2.26673 | 0.0234 | 0.56 |
| Peptidase_M18 | 2.08070 | 0.19883 | 0.08766 | 2.26822 | 0.0233 | 0.56 |
| NADH.G_4Fe.4S_3 | 1.56154 | -0.19853 | 0.08803 | -2.25522 | 0.0241 | 0.569 |
| His_kinase | 2.28043 | 0.19515 | 0.08705 | 2.24179 | 0.025 | 0.581 |
| NMO | 3.56269 | -0.18108 | 0.08206 | -2.20670 | 0.0273 | 0.627 |
| Terminase_4 | 1.43280 | 0.19495 | 0.08856 | 2.20140 | 0.0277 | 0.627 |
| Bac_luciferase | 4.91876 | -0.16502 | 0.07709 | -2.14054 | 0.0323 | 0.722 |
| HisG | 1.65996 | 0.18619 | 0.08852 | 2.10349 | 0.0354 | 0.728 |
| HTH_17 | 1.51178 | 0.18607 | 0.08854 | 2.10148 | 0.0356 | 0.728 |
| IGPD | 1.65996 | 0.18619 | 0.08852 | 2.10349 | 0.0354 | 0.728 |
| Molybdop_Fe4S4 | 1.54617 | -0.18508 | 0.08804 | -2.10224 | 0.0355 | 0.728 |
| Ribosomal_S4e | 1.67540 | 0.17419 | 0.08276 | 2.10488 | 0.0353 | 0.728 |
| SBF | 1.55241 | -0.18588 | 0.08799 | -2.11261 | 0.0346 | 0.728 |
| Ubie_methyltran | 2.80073 | -0.18110 | 0.08551 | -2.11782 | 0.0342 | 0.728 |
| CPDase | 1.36085 | -0.18257 | 0.08754 | -2.08561 | 0.037 | 0.747 |
| Peptidase_M4_C | 2.28903 | 0.18117 | 0.08703 | 2.08157 | 0.0374 | 0.747 |
| GAF | 1.42696 | 0.18185 | 0.08848 | 2.05520 | 0.0399 | 0.752 |
| Methyltransf_15 | 1.35451 | -0.18095 | 0.08772 | -2.06277 | 0.0391 | 0.752 |
| Peptidase_U35 | 1.27255 | 0.18199 | 0.08819 | 2.06367 | 0.039 | 0.752 |
| PFL | 1.64459 | 0.18207 | 0.08853 | 2.05671 | 0.0397 | 0.752 |

| | | | | | | |
|---|---|---|---|---|---|---|
| PRA.PH | 1.66288 | 0.18279 | 0.08851 | 2.06519 | 0.0389 | 0.752 |
| Competence_A | 2.33251 | 0.17716 | 0.08683 | 2.04028 | 0.0413 | 0.771 |
| Acid_phosphat_B | 1.55906 | 0.18008 | 0.08856 | 2.03339 | 0.042 | 0.775 |
| ATP.grasp_3 | 4.92616 | -0.15601 | 0.07695 | -2.02752 | 0.0426 | 0.778 |
| Histidinol_dh | 1.69805 | 0.17892 | 0.08847 | 2.02243 | 0.0431 | 0.779 |
| HSP20 | 1.50674 | -0.17724 | 0.08796 | -2.01494 | 0.0439 | 0.785 |
| PAF.AH_p_II | 1.22877 | -0.17507 | 0.08729 | -2.00564 | 0.0449 | 0.794 |
| BioW | 1.53722 | 0.17722 | 0.08856 | 2.00114 | 0.0454 | 0.794 |
| Condensation | 2.28126 | 0.17159 | 0.08705 | 1.97111 | 0.0487 | 0.836 |
| ICMT | 1.68156 | 0.17431 | 0.08849 | 1.96985 | 0.0489 | 0.836 |

Metrics produced as standard DESeq2 outputs (Love, Huber, & Anders, 2014). **a**: average of the normalized count values, divided by size factors and taken over all samples. **b**: how much the domain presence has changed between the groups studied. Reported on a logarithmic scale (base 2). **c**: standard error for the log2 Fold Change estimate. **d**: Wald statistic value for the Pfam domain tested. **e**: Wald test p-value. **f**: Benjamini-Hochberg adjusted p-value, used for significance cut-off.

### Supplementary Figure 1: Decision tree used for machine learning

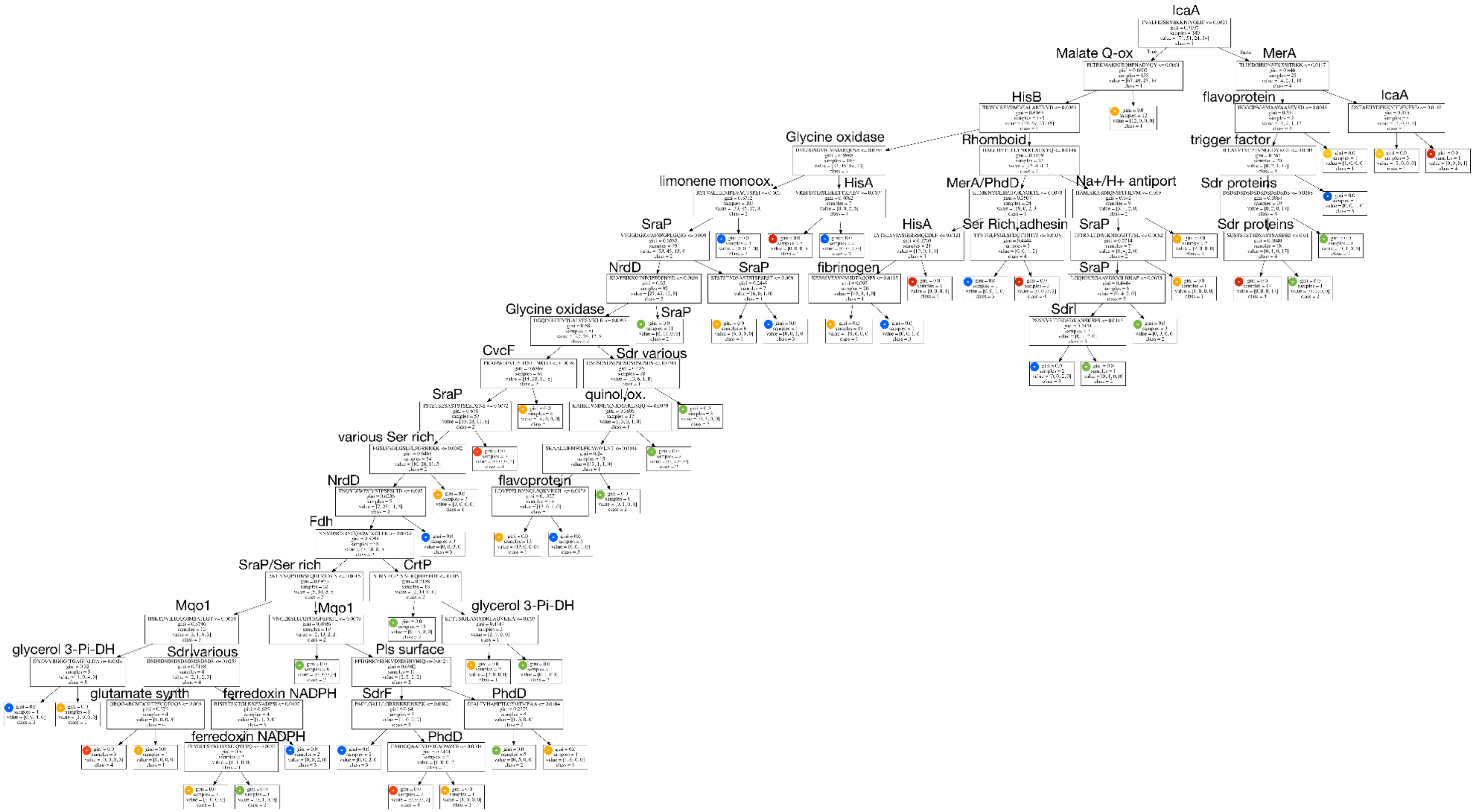

**Supplementary Figure 2: Melt curves and validation graphs for RT-qPCR primers**.

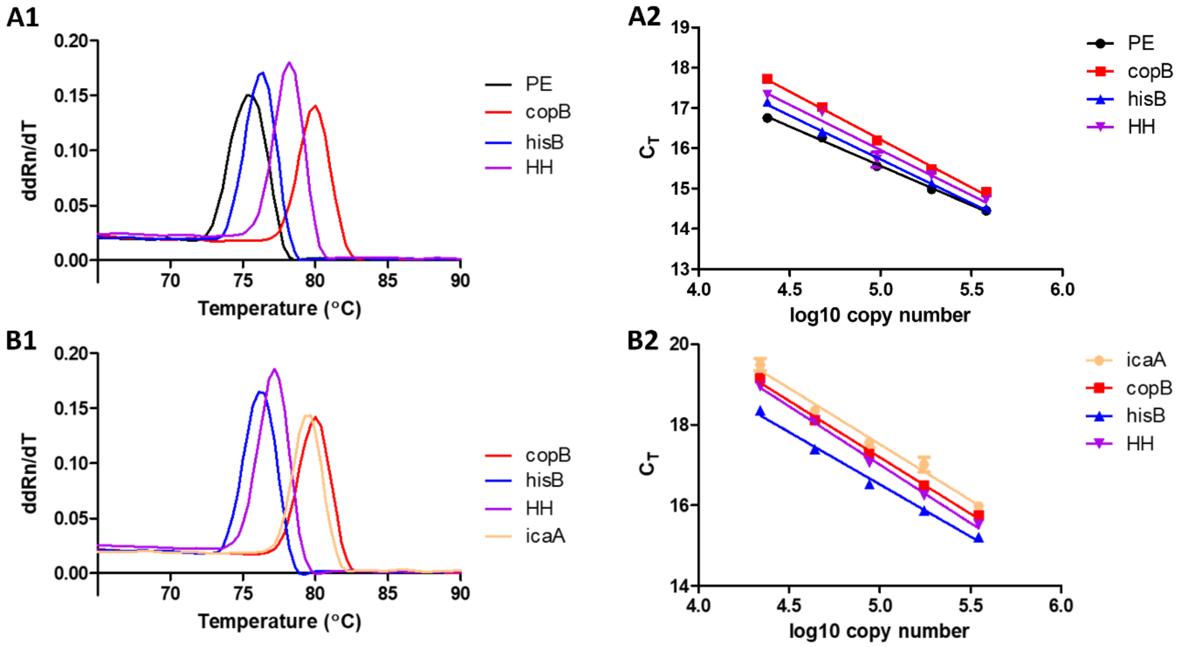


A) *Staphylococcus epidermidis* 846 gDNA used, B) *Staphylococcus epidermidis* RP62A gDNA used. 1) melt curves of RT-qPCR products showing clear single products, 2) CT values obtained across a range of DNA concentrations (each point is the mean of 3 replicates) and line of best fit. Primers were only used where lines of best fit were calculated to have an $R^2$ value above 0.95.
